# LATENT FEEDING BEHAVIORS SUPPORT TROPHIC VERSATILITY IN CICHLIDS

**DOI:** 10.64898/2026.01.21.700746

**Authors:** Khalil T. Russell, Peter C. Wainwright

## Abstract

The relationship between morphology and ecology is mediated by behavior. We explore this relationship by assessing the link between trophic ecology and the use of prey-specific feeding behaviors in a cichlid fish system. Cichlid diversification features repeated transitions between free-moving prey and attached benthic prey, requiring predators to evolve prey-specific approaches to feeding. Using 2000 Hz video, we characterized feeding behavior on an experimental attached benthic prey in seven species of Mesoamerican heroine cichlid spanning three independent transitions to specialized piscivory and two to specialized benthic-feeding ecology. We investigated the effect of feeding ecology on the behavior and kinematics of benthic grazing, a derived, specialized mode of cichlid feeding. Surprisingly, all species readily fed on benthic prey, regardless of their feeding ecology. Nearly all non-benthic species used the same benthic-feeding behaviors as ecological benthic-feeders. Our findings demonstrate an unexpected level of behavioral versatility among cichlid species in exploiting functionally demanding prey outside their typical diets. We propose that this repertoire of latent feeding behaviors supports trophic versatility and facilitates niche diversification. We also show that two benthic-feeding lineages of Neotropical cichlids evolved distinct approaches to benthic feeding, exhibiting the highest and lowest total feeding-strike kinesis, respectively. Together, our findings highlight the importance of behavior in linking morphology and ecology and motivate further study into the diversity and evolutionary context of benthic feeding across the Cichlidae.

**SUMMARY STATEMENT:** We demonstrate that prey-specific feeding behaviors and strike kinematics vary with trophic ecology in heroine cichlids and discuss the potential role of latent feeding behaviors in trophic diversification.

## INTRODUCTION

The Cichlidae are a successful clade of teleosts known for their ecomorphological diversity. Evolutionary transitions between prey types are a prominent feature of cichlid diversification, with extant lineages occupying a wide range of trophic niches (Arbour et al., 2020; McGee et al., 2016; Winemiller et al., 1995). Previous work describes the relationship between functional morphology and trophic ecology among cichlids (Burress, 2016; Hulsey, 2006), highlighting the importance of morphological adaptation in cichlid trophic diversification. However, transitions in prey use require more than morphological changes to the feeding mechanism. For an individual or lineage to successfully transition between prey types, behavior must be sufficiently malleable to support the pursuit and exploitation of non-standard prey (Liem, 1979, 1980). As such, understanding the phylogenetic distribution of behavioral traits in addition to morphology may represent an important step in further understanding trophic diversification.

Evolutionary transitions between prey types may require predators to modify their prey capture strategies. For instance, the capture of evasive, free-moving prey such as fishes often relies on the ability of predatory fish to quickly overtake prey while generating suction by expanding the oral jaws and buccal cavity using complex linkages and skull kinetics (Day et al., 2015; Ferry et al., 2015). In contrast, reliance on sessile, attached, benthic prey may require different behaviors, such as biting, scraping, or tearing, which involve different musculoskeletal mechanisms of the jaws and body (Liem, 1979; Perevolotsky et al., 2020). Cichlids have repeatedly evolved to occupy disparate trophic niches, including specialized piscivory, algivory, molluscivory, and planktivory (Olvera-Ríos et al., 2023). Although it is well known that interspecific differences in diet are often accompanied by interspecific differences in feeding morphology, the relationship between feeding-related behaviors and trophic ecology is less apparent. Species with distinct diets may exhibit distinct feeding behaviors, but it is also possible that specialized lineages retain a diverse suite of behaviors regardless of their present trophic state. Whereas the first scenario would require transitioning lineages to re-evolve prey-type-specific behaviors during trophic niche shifts, the latter might allow individuals to respond to novel niche opportunities with shifts in the use of appropriate feeding behaviors that are part of a versatile feeding repertoire, permitting more rapid transitions between prey.

Biological motions are both powered and constrained by morphology, and it is through the motions of prey capture (e.g., biting, suction-feeding, etc.) that the feeding mechanism acts upon prey. Natural selection, in turn, acts not only upon the feeding mechanism (i.e. trophic functional morphology) but also on how the mechanism is employed during prey capture. Morphological specialization for one mode of prey capture may promote performance on certain prey (e.g., evasive prey capture and evasive prey) but trade-offs are thought to exist that limit performance on other types of prey (e.g., benthic prey capture)(Ferry-Graham & Konow, 2010; Kotrschal, 1988; Westneat, 1994). However, versatility in how species use their morphologies (e.g., behaviors, kinematics) may permit some level of resilience that mitigates the effects of mechanical trade-offs in the feeding system. Understanding this versatility may give insight into the complex relationship between morphology and ecology, and how this relationship changes during lineage diversification.

The heroine cichlids (Heroini) are a Mesoamerican radiation of cichlids that span a range of trophic specialization, with previous work revealing several independent transitions to a wide range of feeding ecologies, including multiple transitions to specialized piscivory, strict herbivory, periphyton scraping, detritivory, and substrate-sifting (Burress & Muñoz, 2021; Říčan et al., 2016; Figure 1). Although these transitions typically follow common evolutionary pathways, there are exceptions to these patterns. For example, although specialized piscivory has predominantly arisen from a more generalized reliance on evasive prey, it also appears to have arisen once from detritivory/molluscivory in the genus *Herichthys/Nosferatu* (Perez-Miranda et al., 2019; Říčan et al., 2008, 2016). Such a transition between disparate prey types (benthic prey to evasive prey) highlights the potential importance of trophic versatility in Heroine cichlids in facilitating niche divergence and diversification. While some studies have begun to describe the distribution of behavioral and kinematic variation among cichlids more broadly (Higham et al., 2007; Liem, 1978, 1980; Martinez et al., 2024), their ecomorphological diversity makes them an excellent system for exploring variation in feeding strategies as they relate to feeding ecology.

**Figure 1:**
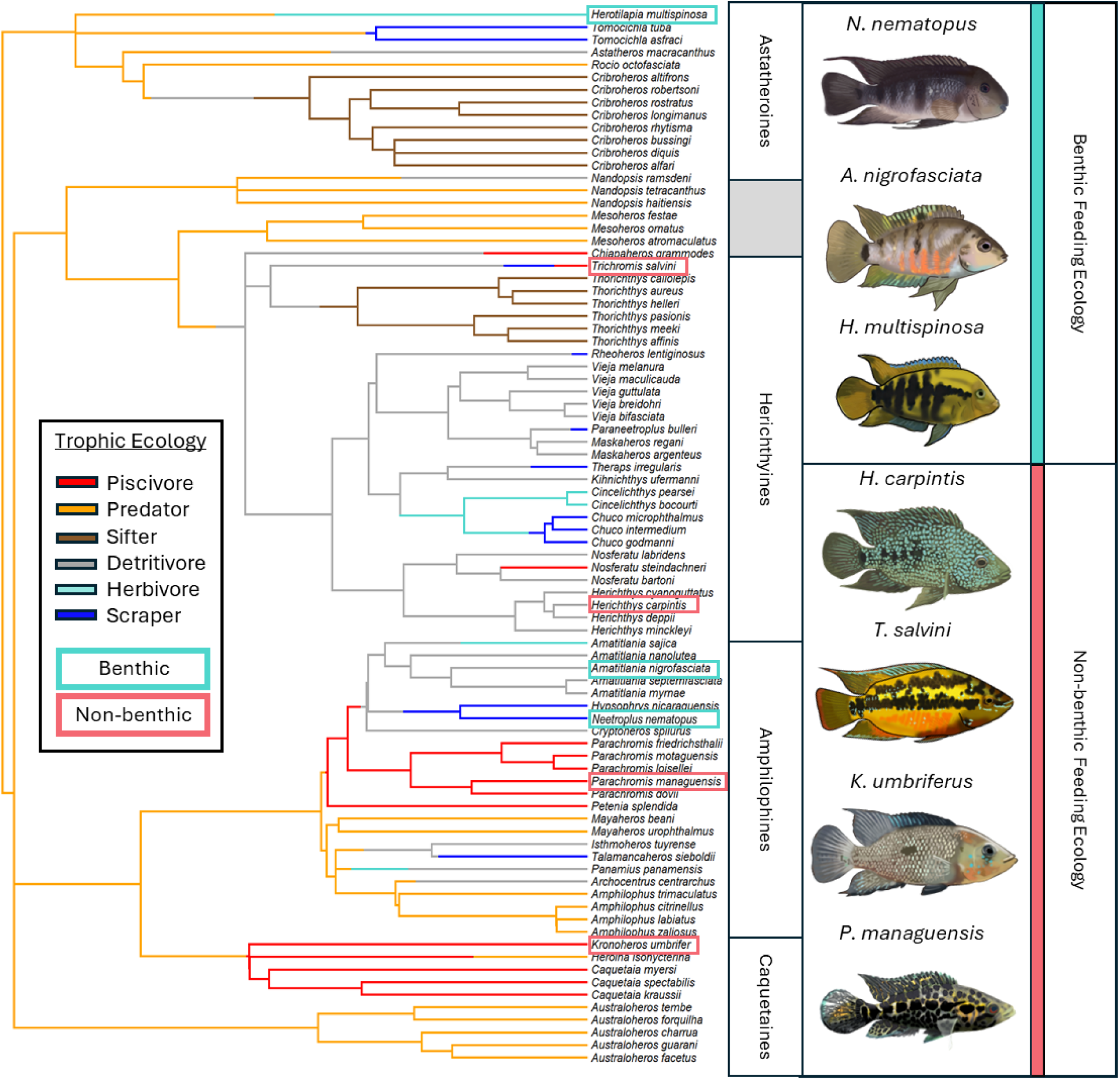
Phylogeny of the Heroini overlaid with an ancestral-state reconstruction of trophic ecology. Phylogeny is from McGee et al. (2020), trophic ecology from Říčan et al. (2016). Species assessed in the present study are in boxes: cyan—benthic ecology; red—non-benthic ecology. Fish illustrations by Khalil Russell.

In the present study, we aimed to assess and describe the relationship between trophic ecology and benthic-feeding kinematics and behaviors in heroine cichlids. Previous work suggests that the ancestral trophic condition for cichlids is generalized invertivory, and that specialized benthic-feeding, such as periphyton scraping or strict herbivory, is a derived mode of prey capture among heroine cichlids (Říčan et al., 2016). It is yet unclear how cichlids primarily reliant on evasive prey feed on benthic prey, or whether they are willing to do so at all. Understanding the distribution of specialized behaviors related to benthic feeding may help clarify how lineages transition between reliance on primarily midwater evasive prey and attached benthic prey. We first determined the extent to which the fundamental ability and willingness to feed on attached, benthic prey is distributed across ecologically diverse heroine species. To address this, we presented an experimental benthic food to seven species spanning a range of ecological reliance on evasive prey (0-92%). We then sought to characterize benthic-feeding behaviors and kinematics to determine the effect of trophic ecology on benthic-feeding. Our expectation was that willingness to feed on the experimental, benthic food would be reduced in species that normally feed on elusive, mobile prey. We further predicted that kinematics would differ among species, with benthic feeding ecology being associated with distinct kinematic and behavioral traits.

## MATERIALS & METHODS

### Species Selection

Among the seven species chosen, three rely on evasive prey (Hulsey & García De León, 2005) for one percent or less of their diets: *Amatitlania nigrofasciata, Neetroplus nematopus,* and *Herotilapia multispinosa*. For two species, evasive prey form 25% and 48% of the diet, respectively: *Herichthys carpintis* (Perez-Miranda et al., 2019) and *Trichromis salvini* (Hulsey & García De León, 2005). Evasive prey forms 90% or more of the diet of the final two species: *Kronoheros umbriferus* (K. Winemiller, personal communication, July 26, 2025) and *Parachromis managuensis* (Hulsey & García De León, 2005). Species were also chosen to represent different lineages within the Heroine cichlid radiation (Table 1). Three species belong to the amphilophine clade (*P. managuensis, N. nematopus, A. nigrofasciata*), two belong to the hericthyine clade (*T. salvini, H. carpintis*), one to the astatheroines (*H. multispinosa*), and one to the caquetaines (*K. umbriferus*). Our study contains two independent transitions to specialized benthic-feeding among Heroine cichlids (Figure 1): (the amphilophine species and the astatheroine species), as well as three distinct benthic-feeding ecologies (detritivory/picking—*A. nigrofasciata*, periphyton scraping—*N. nematopus*, algivory*—H. multispinosa;* see Table 1 for literature cited). Among our non-benthic-feeders, we represent three independent transitions to specialized piscivory across three subclades of Heroine cichlids: *K. umbriferus*—caquetaine, *P. managuensis*—amphilophine, and *T. salvini*—herichthyine (Říčan et al., 2016).

**Table 1.**
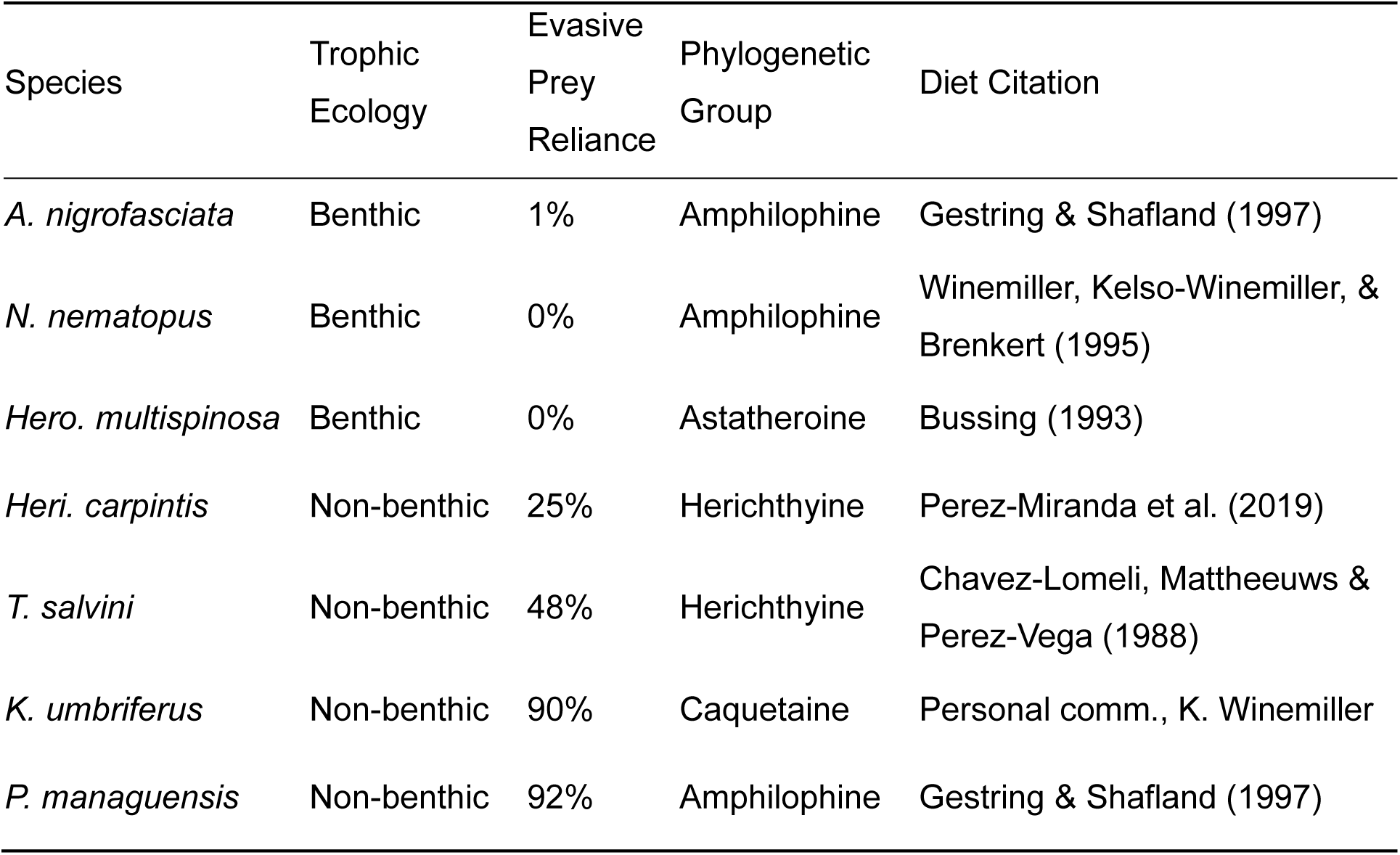
Trophic ecology and taxonomy of study species. Reliance on evasive prey primarily from Hulsey & García De León (2005). *Kronoheros umbriferus* ecology was estimated from personal communication with Kirk Winemiller and morphological characterization by Říčan et al. (2016).

### High-speed Filming

Species were housed and filmed singly at room temperature in 109-liter aquaria, and for about two weeks prior to filming were fed primarily with the experimental food, which consisted of a mixture of gelatin and ground fish food (Hikari Cichlid Staple) set on a microscope slide. The food mixture was added to cover an area of approximately 2cm^3^ (1cm × 2cm) at a mean thickness of <1mm. This experimental food plate was designed to elicit a benthic-feeding response, where fish must physically remove attached food via contact between the jaws and food as opposed to sucking it from the water column.

We filmed 33 individuals across the seven species (minimum = 4, maximum = 6 individuals). Once transferred to filming aquaria, subjects were exposed to the diet treatment under different lighting in an effort to acclimatize them to the filming conditions. Each fish was filmed over a period of one to several weeks until 20 acceptable sequences were obtained, with the most cooperative subjects completing filming in a single day (i.e., all films were collected on the individual in a single filming session). The filming setup consisted of the benthic food plate attached by a rubber band to the top of an acrylic block.

Subjects were filmed at 2000 frames per second using a digital Photron Fastcam Mini camera fitted with an AF Nikkor 50mm f/1.8D lens. Each film captured a single benthic-feeding strike, which begins with the start of jaw protrusion, includes contact between one or both jaws and the substrate, and ends with complete jaw retraction into the skull. Occasionally, subjects would engage in multiple open-close cycles of the jaws between initial jaw protrusion and complete jaw retraction—these films were not used. Subjects were filmed 10 times in an approximately lateral view, where the anterior-posterior axis of the fish was parallel to the camera lens, as well as 10 times in a dorsal view, where the anterior-posterior axis was perpendicular to the camera. Sequences were illuminated with a bank of LED lights. All experimental procedures and animal maintenance practices were approved by the UC Davis IACUC committee under protocol #23818.

We extracted six frames from each lateral film with the software ImageJ (Schneider, Rasband, & Eliceiri, 2012) based on events during the feeding strike shared across all species observed (Figure 2). The first frame (F0) captured the start of jaw protrusion in the upper jaw; F1 captured the initial moment of contact between the lower jaw and the food plate; F2 captured the point of peak upper jaw protrusion, and corresponds with the point of maximal contact with the substrate in all species observed; F3 captures the moment of maximal jaw closure prior to either jaw retraction or loss of contact with the substrate. Frames 4 and 5 are alternate frames, where each strike sequence exhibits either F4 or F5, but not both (Figure 2). Following jaw closure, jaw retraction either resulted in the movement of the head toward the relatively stationary tips of the jaws (F4), which would maintain their position against the substrate, or the movement of the jaws toward the relatively stationary head (F5). The final frame, F6, was the moment when the jaws were fully retracted into the skull and no longer in contact with the substrate.

**Figure 2:**
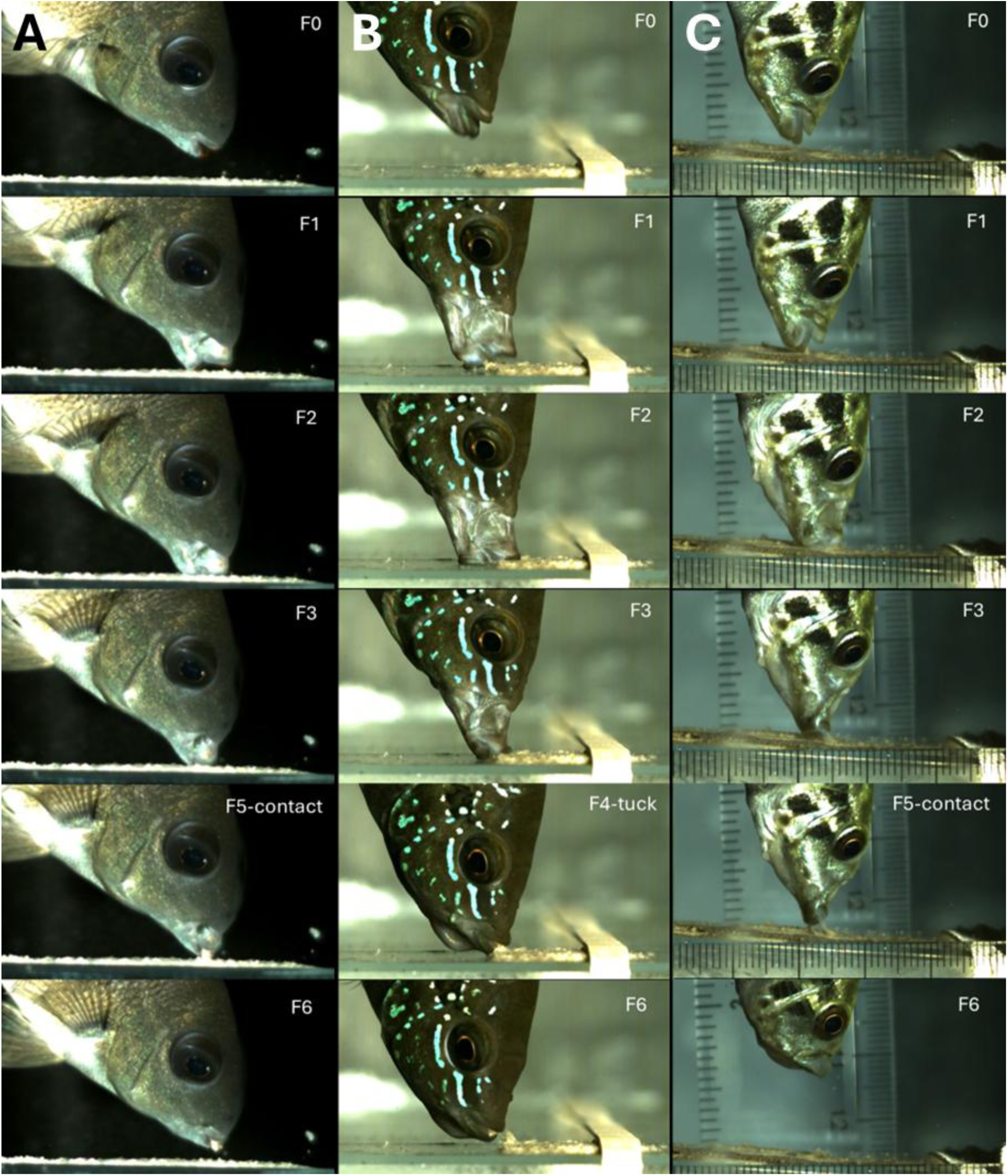
Benthic-feeding strike sequences from three cichlid species: A) *Neetroplus nematopus*; B) *Trichromis salvini;* C) *Parachromis managuensis*. Frame 0—start of jaw protrusion; Frame 1—initial contact with substrate; Frame 2—peak protrusion; Frame 3—jaw closure prior to retraction; Frame 4 (alternating)—jaw retraction with substrate contact; Frame 5 (alternating)—loss of substrate contact; Frame 6—jaws mostly retracted.

Each strike sequence consisted of six total frames: F0, F1, F2, F3, F4/F5, and F6. We used tpsDig (Rohlf, 2006) to place seven landmarks on each frame across each film (*n* = 300). Each landmark was placed on a homologous anatomical position across each subject in each frame (Figure 3).

**Figure 3:**
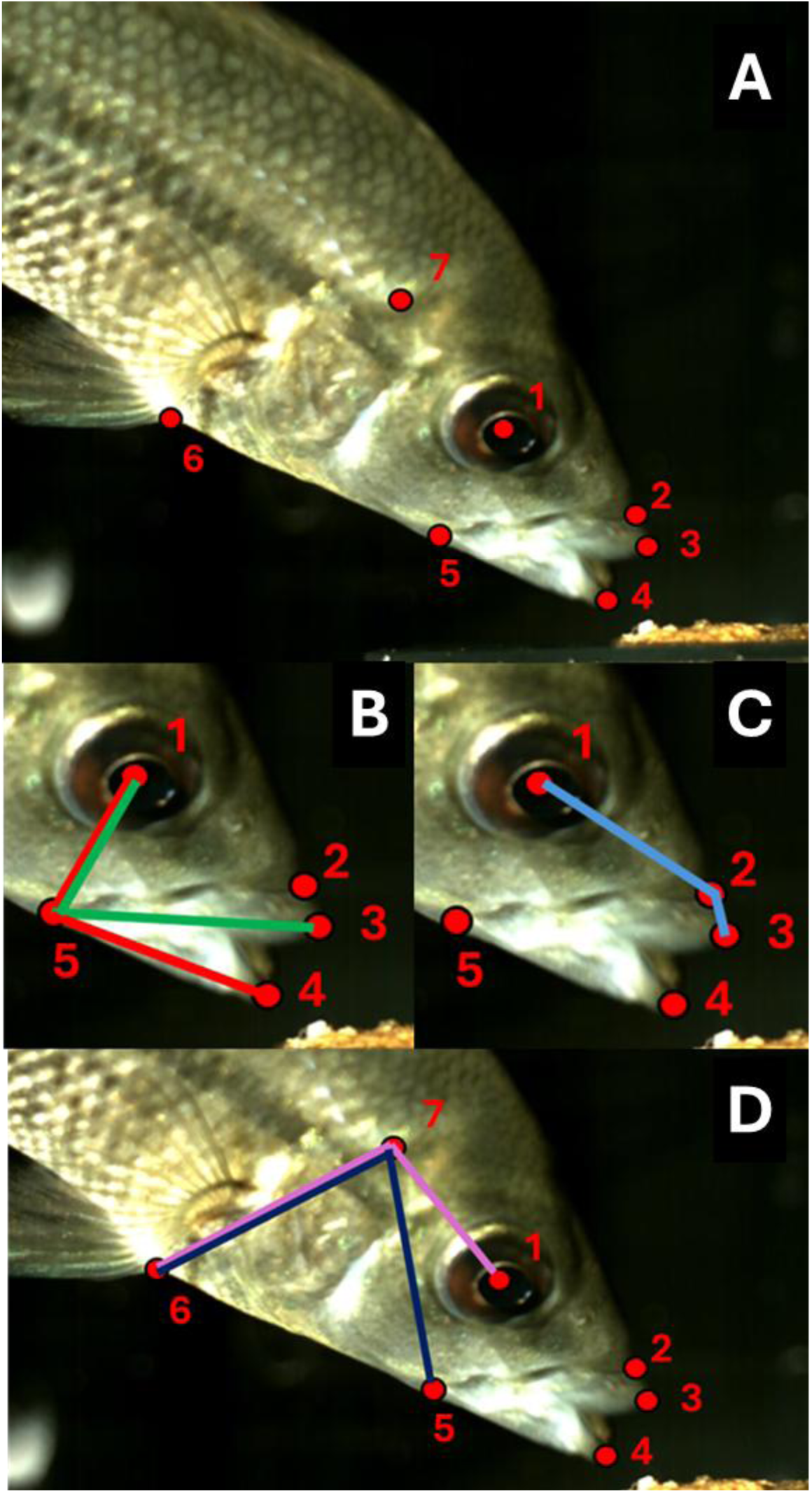
A - Landmarking scheme for kinematic data collection. Seven landmarks were placed on each frame for each film: (1) center of orbit; (2) tip of snout; (3) tip of upper jaw; (4) tip of lower jaw; (5) lower-jaw joint; (6) pelvic fin insertion; (7) point of cranial rotation. Landmarks were used to calculate the following angles: B (green) – upper-jaw angle; B (red) – lower-jaw depression angle; C – upper-jaw scrape angle; D (magenta) – cranial rotation; D (indigo) – pelvic expansion.

### Variables – Kinematic motions & benthic-feeding behaviors

Eight variables were extracted from the lateral films: five kinematic motion variables and three behavioral variables. We measured both internal motion—that is, the relative motion of one body part versus another—and external motion—the motion of the fish or its parts relative to the environment. We emphasize this distinction by referring to the former as kinematics and the latter as behaviors. The following kinematic variables were calculated from the coordinate data on each landmarked lateral film frame using RStudio (RStudio Team, 2019): upper-jaw angle (angle from landmarks 1-5-3), lower-jaw depression angle (1-5-4), upper-jaw scrape angle (1-2-3), cranial rotation angle (6-7-1), and pelvic expansion angle (6-7-5; Figure 3). We took the range of each kinematic variable through the strike for further analysis.

A total of four behavioral variables were collected from films: strike variability, jaw contact rate, body pitch angle, and head-flick rate. *Strike variability* was defined as the proportion of lateral strike sequences for an individual with an F4 frame, as opposed to an F5. Jaw contact was measured by counting the number of jaws in contact with the experimental food for F2 from each lateral strike sequence. *Jaw contact rate*, in turn, was the proportion of films in which both the upper and lower jaw contacted the food at this time, as opposed to partial jaw-contact, where only the lower jaw contacts the food at peak protrusion (Figure 4). *Lateral body pitch angle* was the angle between a line through the center of the caudal fin insertion and the lower jaw point of contact with the experimental food, and the horizontal surface of the experimental food. *Head-flick rate* was calculated as the proportion of dorsal film strikes that included a lateral head-flick, a behavior that occurred at the end of a bite in which the fish quickly moved its head laterally.

**Figure 4:**
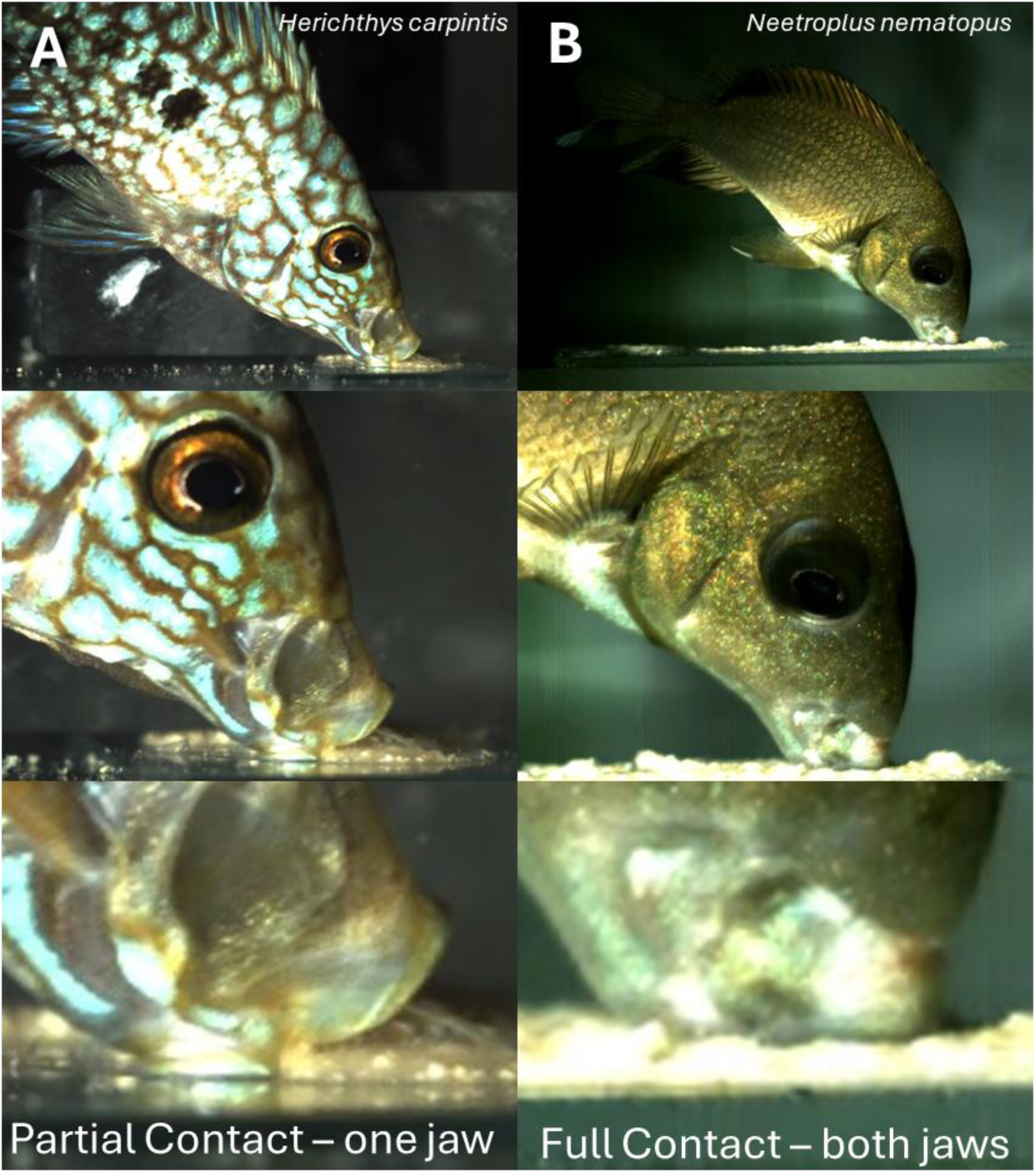
Frame 2 from 2000 Hz films of *Herichthys carpintis* (A) and *Neetroplus nematopus* (B) feeding on experimental food. A-partial contact between the jaws and the substrate. B-complete contact between the jaws and the substrate.

### Statistical Analysis

To summarize the patterns of kinematic and behavioral variation among species, we performed three sets of principal components analyses on the correlation matrices of measured variables. The first PCA included only the range of excursions of the five kinematic variables, the second included only the four behavioral variables, and the third combined all nine variables in a single analysis. Principal components analyses were performed on individual means, and species means were calculated by averaging principal component scores across individuals. To test for the effect of trophic ecology on feeding movements, we also performed phylogenetic least squares regressions (PGLS) with the phylogeny from (McGee et al., 2020) on the relationship between PC1 of the combined PCA and percent evasive prey in the diet (species means), as well as the relationship between body pitch angle and rates of complete jaw-contact (PGLS of individual means).

## RESULTS

We observed a general benthic-feeding strike pattern across all seven species. Feeding strikes were preceded by a body orientation phase, where the fish positioned itself relative to the substrate before jaw protrusion (Figure 2—F0). After this, lower jaw depression and upper jaw protrusion led to either partial (one jaw) or complete (both jaws) contact between the jaws and the substrate, where initial contact was exclusively made with the lower jaw (Figure 2—F1). Partial contact was virtually always made between the lower jaw and substrate, and complete contact between both jaws and the substrate coincided with the point of maximum gape and jaw protrusion (Figure 2—F2; Figure 4). Jaw-closure occurred next, where either one or both jaws moved to close the distance between the tips of the upper and lower jaws. In strike sequences with dual jaw-contact, one of the jaws served as an apparent anchor, unmoving in its point of contact with the substrate (Figure 2—F3). This often resulted in the removal of benthic food via scraping between the moving jaw and the substrate. In strike sequences with single jaw-contact, the movement of the non-anchor jaw toward the anchor jaw resulted in jaw closure. Following jaw closure, if the jaws were closed around a bit of food that was firmly attached to the substrate, jaw retraction resulted in the movement of the head and body downward toward the substrate (Figure 2—F4). This was often, but not always, followed by a lateral head flick, where the subject jerked its head to one side or the other. When jaw-closure occurred without a grip on firmly attached food, the processes of jaw-closure and retraction virtually always resulted in loss of contact between the fish and the substrate (Figure 2—F5-contact).

**Figure 5:**
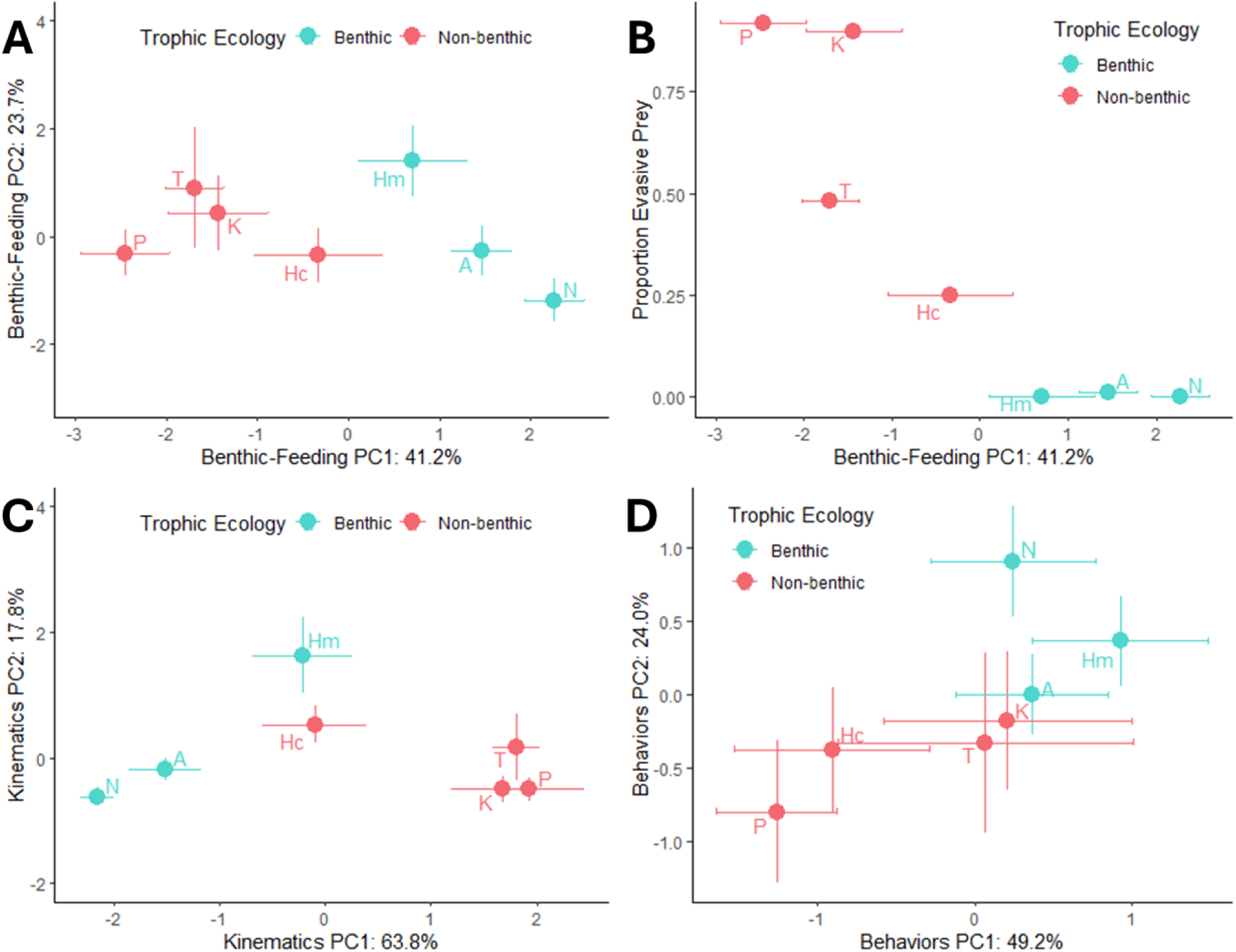
A) Principal components analysis of benthic-feeding behavioral and kinematic variables combined. B) PC1 from combined-variable PCA versus the proportion of the diet composed of evasive prey. C) Principal components analysis of only kinematic variables. D) Principal components analysis of only behavioral variables. Species means and standard error are plotted for all graphs. Species codes: A—*A. nigrofasciata* (n=6), Hc—*H. carpintis* (*n* = 4), Hm—*H. multispinosa* (*n* = 4), K—*K. umbriferus* (*n* = 5), N—*N. nematopus* (*n* = 6), P—*P.. managuensis* (*n* = 4), T—*T. salvini* (*n* = 4).

We observed two distinct patterns of benthic-feeding strikes (Figure 2), where the sequence employed was partially a function of the grip between the subject’s jaws and the substrate. The proportion of strikes that showed the F4 pattern was a function of trophic ecology, as the species with the greatest reliance on evasive prey (*Parachromis managuensis*, 92%; Table 1) was only ever observed using a single strike pattern (0% strike variability; Table 2; Figure 2—A,C). In contrast, the greatest strike variability (45%; Table 2) was observed for one of the species tied for the least reliance on evasive prey in our dataset (*Herotilapia multispinosa*, 0%; Table 1).

**Table 2.**
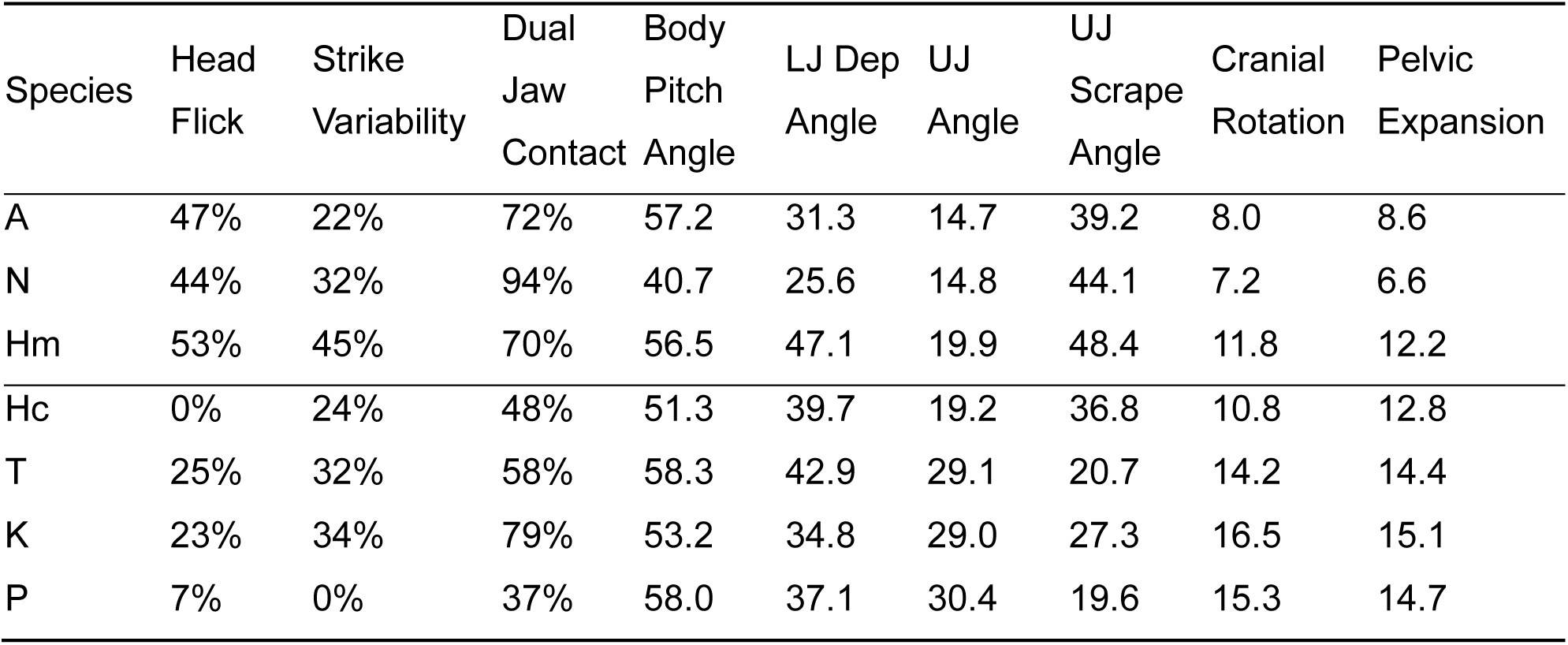
Kinematic and behavioral means across examined species. Method of calculating the first three variables is described in the text. The five variables to the right are all the range of each angle, in degrees, from onset of the strike to maximum excursion. Species ecology: benthic—A, N, Hm; non-benthic—Hc, T, K, P. Species codes: A—*A. nigrofasciata* (n=6), Hc—*H. carpintis* (*n* = 4), Hm—*H. multispinosa* (*n* = 4), K—*K. umbriferus* (*n* = 5), N—*N. nematopus* (*n* = 6), P—*P.. managuensis* (*n* = 4), T—*T. salvini* (*n* = 4).

| Species | Head Flick | Strike Variability | Dual Jaw Contact | Body Pitch Angle | LJ Dep Angle | UJ Angle | UJ Scrape Angle | Cranial Rotation | Pelvic Expansion |
| --- | --- | --- | --- | --- | --- | --- | --- | --- | --- |
| A | 47% | 22% | 72% | 57.2 | 31.3 | 14.7 | 39.2 | 8.0 | 8.6 |
| N | 44% | 32% | 94% | 40.7 | 25.6 | 14.8 | 44.1 | 7.2 | 6.6 |
| Hm | 53% | 45% | 70% | 56.5 | 47.1 | 19.9 | 48.4 | 11.8 | 12.2 |
| Hc | 0% | 24% | 48% | 51.3 | 39.7 | 19.2 | 36.8 | 10.8 | 12.8 |
| T | 25% | 32% | 58% | 58.3 | 42.9 | 29.1 | 20.7 | 14.2 | 14.4 |
| K | 23% | 34% | 79% | 53.2 | 34.8 | 29.0 | 27.3 | 16.5 | 15.1 |
| P | 7% | 0% | 37% | 58.0 | 37.1 | 30.4 | 19.6 | 15.3 | 14.7 |

There was a clear relationship in which species with a benthic-feeding ecology were more willing to feed on the experimental food preparation. On average, piscivores (*Ts, Pm, Ku)* took 3.58 sessions, whereas *Neetroplus nematopus* (scraper) took 1.3 sessions to film, with 4 individuals completing filming in one session each (20 films), and *Herotilapia multispinosa* (herbivore) took 2.5 sessions on average. *Amatitlania nigrofasciata* took the highest average number of sessions to film.

The principal components analysis performed on all nine variables (behaviors + motions) resulted in a first PC which captured 41.2% of total variation, and a second principal component which captured 23.7% of the variation (Figure 5a). Positive values on the first principal component are associated with greater ranges of motion of the upper-jaw scrape angle, higher rates of lateral head flicks, and greater jaw-contact with the substrate. Negative PC1 values are associated with greater ranges of motion of the upper-jaw angle, cranial rotation angle, and pelvic expansion angle. Principal component 2 is positively correlated with greater strike variability, lower-jaw depression angle ranges, and steeper body pitch angles.

We regressed the first principal component against the proportion of evasive prey in the diet and observed a negative relationship between PC1 values and reliance on evasive prey (Figure 5b). We performed this analysis twice—once including *Kronoheros umbriferus*, whose reliance on evasive prey was estimated (personal communication, K. Winemiller), and once using only species whose reliance on evasive prey was quantified in the literature (all but *K. umbriferus*). Both analyses yielded significance in the relationship between PC1 and proportion of evasive prey in the diet (p = 0.04, p < 0.0001, respectively).

A principal components analysis performed with the five kinematic motion variables resulted in the primary axis of variation capturing 63.8% of the total variation (Figure 5c). Here, positive values of PC1 are associated with greater ranges of motion in the upper-jaw angle, cranial rotation, and pelvic expansion, and smaller ranges of motion in the upper-jaw scrape angle. The second principal component captured 17.8% of the variation and is positively correlated with the ranges of motion in the lower-jaw depression angle and upper-jaw scrape angle (Figure 5).

The PCA of the four benthic-feeding behaviors resulted in a primary axis of variation which captured 49.2% of the total variation, and a secondary axis which describes 24.0% (Figure 5d). The first principal component is positively correlated with all four variables: head-flick rate, strike variability, jaw-contact rate, and body pitch angle. The second principal component is positively correlated with head flicks and strike variability and negatively correlated with contact rate and body pitch angle. Along PC1, there is almost complete overlap between benthic- and non-benthic-feeders, with only the negative-most values of PC1 reserved for non-benthic-feeders only. Across most of the PCA, principal component 2 shows some separation between benthic- and non-benthic-feeders, with positive values largely dominated by benthic-feeders, and negative values largely occupied by non-benthic-feeders. However, for the lowest values of PC1, there are non-benthic-feeders which overlap with benthic-feeders along PC2.

We calculated a measure of total kinesis by summing the mean ranges of angular motion for each species (Figure 6). The highest total kinesis was observed in *H. multispinosa*, and the lowest in *N. nematopus* and *Amatitlania nigrofasciata*. The three benthic-feeding species (*A. nigrofasciata*, *N. nematopus*, *H. multispinosa*) exhibited greater range in total kinesis than the four non-benthic-feeders (*Herichthys carpintis*, *Kronoheros umbriferus*, *Trichromis salvini*, and *P. managuensis*). This pattern is driven in part by the range of motion for lower-jaw depression, which is highest among the seven species in *H. multispinosa*, and lowest for *A. nigrofasciata* and *N. nematopus*. We also observe *N. nematopus* and *A. nigrofasciata* to have the lowest ranges of cranial rotation and upper-jaw angle, though this is not true for *H. multispinosa*. We found consistently greater ranges of motion for the upper-jaw scrape angle among benthic-feeders versus non-benthic-feeders. In contrast, non-benthic-feeders exhibit consistently greater ranges of motion in pelvic expansion compared to benthic-feeders.

**Figure 6:**
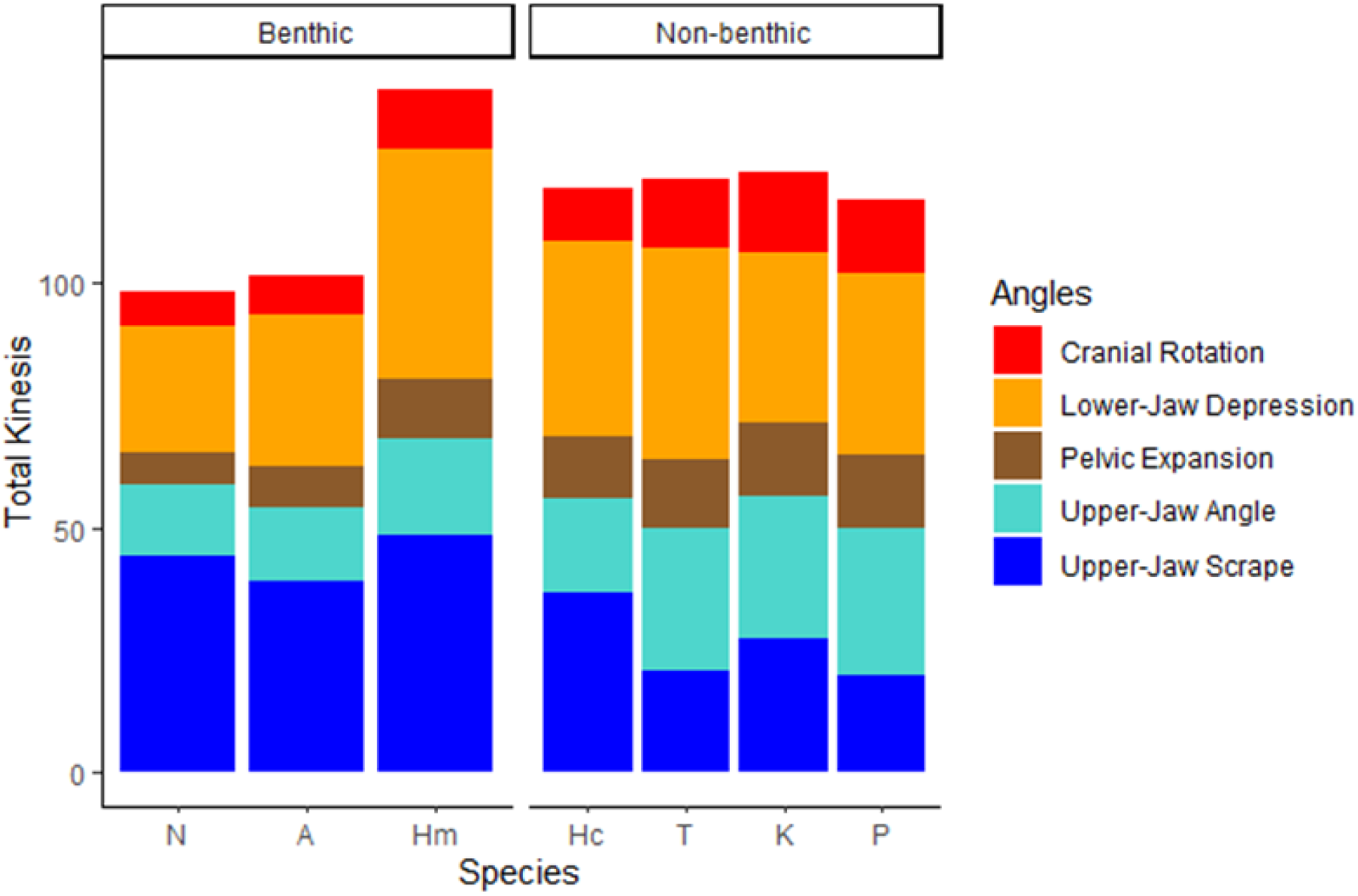
Total kinesis across seven cichlid species (species means). Benthic-feeding ecology is associated with greater movement in the upper-jaw scrape angle and less pelvic expansion, cranial rotation, and upper-jaw angle excursion. Benthic-feeding cichlids from two independent transitions to benthic-feeding ecology exhibit the highest (Hm) and lowest (A, N) total kinesis across the species examined. Hm exhibits higher lower-jaw depression compared to the other benthic-feeders. Species codes: A—*A. nigrofasciata* (n=6), Hc—*H. carpintis* (*n* = 4), Hm—*H. multispinosa* (*n* = 4), K—*K. umbriferus* (*n* = 5), N—*N. nematopus* (*n* = 6), P—*P.. managuensis* (*n* = 4), T—*T. salvini* (*n* = 4).

When assessing variables individually, we observe that head-flick rates are consistently higher among benthic-feeders (44-53%) compared to non-benthic-feeders (0-25%, Table 2). Rates of complete jaw-contact are more variable among non-benthic-feeders (37-79%), whereas they are consistently high among benthic-feeding cichlids (70-94%). The lowest strike variability was observed in *Parachromis managuensis* (0%), the species with the greatest reliance on evasive prey as a proportion of the diet (92%). In contrast, *H. multispinosa* exhibited the greatest strike variability (45%), with the remaining five species exhibiting substantial overlap (22-34%). Mean body pitch angle ranged from 51-58 degrees for all species except *Neetroplus nematopus*, with a mean of 40.7 degrees. The phylogenetic least squares regression yielded a positive significant (R^2^ = 0.584, p = 0.001) relationship between individual-mean body-pitch angles and rates of complete jaw-contact (Figure 7).

**Figure 7:**
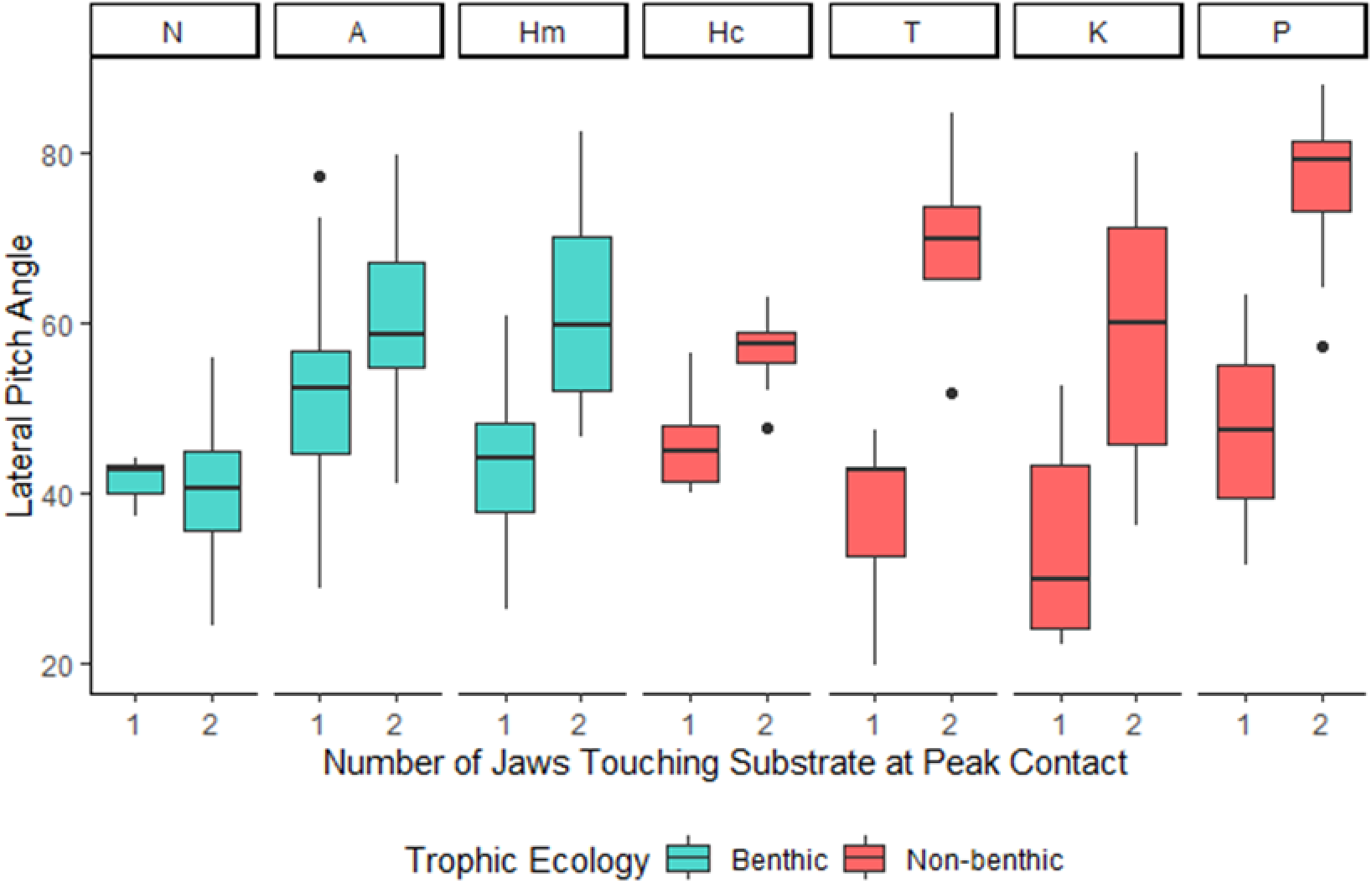
Relationship between jaw-contact and lateral pitch angle in seven neotropical cichlid species. Jaw-contact is the number of jaws (1—lower jaw; 2—upper- and lower-jaws) in contact with the substrate at peak jaw-contact (peak protrusion, frame 2). There is a general relationship between jaw-contact and lateral pitch angle, where full jaw-contact is correlated with steeper pitch angles compared to partial jaw-contact. However, N maintains complete jaw-contact while showing relatively low pitch angles. Species codes: A—*A. nigrofasciata* (n=6), Hc—*H. carpintis* (*n* = 4), Hm—*H. multispinosa* (*n* = 4), K—*K. umbriferus* (*n* = 5), N—*N. nematopus* (*n* = 6), P—*P.. managuensis* (*n* = 4), T—*T. salvini* (*n* = 4).

## DISCUSSION

All species in our study fed on benthic prey regardless of feeding ecology (Supplemental Video 1), suggesting that the fundamental ability and willingness to feed on attached prey is widespread, potentially ubiquitous, among the Heroini. Additionally, all species shared a similar general strike pattern, as well as a set of shared behaviors related to benthic feeding. Nevertheless, feeding ecology had a strong effect on behavior and strike kinematics across the species, supporting the idea that the link between morphology and ecology in the feeding mechanism of fishes is mediated by a link between ecology, kinematics, and feeding-related behaviors.

### Latent benthic-feeding behaviors may promote trophic versatility and diversification

Heroine cichlid diversification commonly features transitions between reliance on evasive mobile prey and attached benthic prey (Burress & Muñoz, 2021; Říčan et al., 2016), and latent benthic-feeding behaviors may play an important role in these transitions. We observed that cichlids primarily reliant on non-benthic, evasive prey can engage in a range of specialized behaviors related to benthic feeding. The willingness of relatively specialized predators to engage in benthic feeding behaviors may ease transitions between prey types with opposing functional properties by removing the constraining necessity of first evolving novel prey-capture behaviors.

Behavioral versatility may also be key to intraspecific variation in trophic ecology within and between cichlid populations. Previous work demonstrates that cichlid populations can vary substantially in their diet composition (Pease et al., 2018). For example, in habitats characterized by seasonal fluctuation in prey availability and prey community composition, latent prey-specific feeding behaviors may allow fishes that are relatively specialized in one season to feed more broadly during another, when their preferred prey may be less abundant (da Costa & Soares, 2015; Duarte et al., 2022). Pronounced niche divergence has also been documented between native and non-native populations of neotropical cichlids (Bergmann & Motta, 2005). The wide success of heroine cichlids during species introductions may similarly be bolstered by their trophic adaptability, especially as the species composition of recipient communities may differ substantially from that of the source community (Comte et al., 2016).

### Adaptations for benthic-feeding distinguish benthic-feeders from evasive-prey specialists

Benthic-feeding cichlids from two independent transitions to specialized benthic feeding, one in the amphilophines and one in the astatheroines, shared high rates of dual jaw contact, greater reliance on lateral head flicks, and greater ranges of excursion in the upper-jaw scrape angle compared to cichlids with non-benthic ecology. From this, it appears that mobility in the upper jaw may represent a key adaptation for benthic-feeding among the Heroini, as it may permit greater rates of dual jaw contact between the jaws and substrate during feeding. The ability to consistently engage both jaws with the substrate may allow benthic-feeding cichlids to more effectively scrape material compared to single jaw contact, resulting in potential benefits to feeding performance on attached prey.

Upper-jaw mobility might also be crucial for effectively using the lateral head flick when removing food. During feeding strikes where only one jaw touches the substrate, the advantage of the lateral head flick for benthic prey capture is unclear because it almost immediately causes a loss of contact between the lower jaw and the substrate. However, when both jaws tightly grip the prey during a strike with dual jaw contact, the lateral head flick can boost the total force the predator applies to the prey by recruiting the muscles that power the head flick for prey capture (Perevolotsky et al., 2020). Upper-jaw mobility may help engage both jaws better during the feeding strike, which could, in turn, improve the effectiveness of the lateral head flick.

Ecological reliance on highly evasive prey is known to be associated with highly kinetic suction-feeding strikes in cichlids (Martinez et al., 2018). We observed consistently elevated cranial rotation, pelvic expansion, and excursion of the upper-jaw angle among non-benthic feeders compared to benthic-feeding cichlids. Kinematic patterns among evasive prey specialists may reflect evolutionary pressures toward high kinesis in the feeding strike, which helps produce rapid expansion of the oral and buccal cavities (Day et al., 2015; Ferry et al., 2015; Westneat & Olsen, 2015). In this way, observed differences between benthic- and non-benthic feeding cichlids in benthic-feeding kinematics may reflect a tradeoff between highly kinetic suction-feeding strikes and highly dexterous benthic-feeding strikes, as has been postulated previously (Burress & Muñoz, 2023; Corn et al., 2021; Westneat, 1994). In the context of previous work, our results suggest that we might observe an associated tradeoff in feeding performance. If alignment between prey type and prey-specific kinematic motions confers advantages in feeding performance, we might expect a tradeoff between highly kinetic and highly dexterous feeding motions to correspond with a tradeoff in feeding performance between evasive and benthic prey.

### Variation among specialized benthic-feeding cichlids

Despite broad patterns separating benthic- and non-benthic-feeding cichlids, our study also revealed differences among benthic-feeders. For example, our measure of total kinesis, which sums across all measured ranges of angular excursion for each species, showed that benthic-feeding cichlids exhibit both the lowest (*Neetroplus nematopus*, *Amatitlania nigrofasciata*) and highest (*Herotilapia multispinosa*) total kinesis, contrasting with comparatively moderate values of total kinesis among non-benthic-feeding cichlids. The difference between the amphilophine and astatheroine benthic-feeders is largely explained by pronounced differences in the range of excursion of the lower-jaw depression angle, where *H. multispinosa* exhibited the greatest degree of lower-jaw depression across the species examined, and the amphilophine cichlids exhibited the least.

Compared to the amphilophine benthic-feeding species, *H. multispinosa* more closely resembles the non-benthic species in kinematics, but not behaviors. When considering behaviors without kinematics, *H. multispinosa* exhibits the highest values along PC1 of all species, with the amphilophine benthic feeders exhibiting greater similarity to non-benthic feeders along PC1. Conversely, *H. multispinosa* is near the center along PC1 on the principal components analysis of kinematic angles of excursion, hardly distinguishable from *Herichthys carpintis* along the primary axis of variation. Here, the amphilophine cichlids represent the lowest values across the seven species. These results may suggest that the two groups of benthic feeders may rely on separate strategies of benthic feeding specialization, with the amphilophine cichlids exhibiting greater kinematic, but less behavioral distinction from non-benthic feeding cichlids compared to *H. multispinosa*. Our results suggest that different lineages may evolve along different paths (i.e., kinematics vs behaviors) to achieve similar outcomes when all variables are considered in combination. Further work is needed to evaluate this possibility.

We also observed that the number of jaws contacting the substrate during the feeding strike correlated with the lateral pitch angle between the body and the substrate, where increasing body pitch angles yielded higher rates of dual jaw contact compared to single jaw contact. This relationship between pitch angle and jaw contact was observed as both a within-species trend, where the pitch angle of a given strike related to whether the strike had single- or dual-jaw contact, but also as an interspecific pattern, where the average body pitch angle and rate of dual jaw contact were correlated across species. However, *Neetroplus nematopus* maintained the highest rates of jaw contact (95%, species mean) despite the lowest body pitch angles (40.7 degrees, species mean; Table 2).

The unique relationship between body pitch angle and complete jaw-contact in *N. nematopus* may reflect an adaptation for benthic feeding efficiency compared to other species, including other benthic feeders. Between feeding strikes, we observed that many subjects spent time reorienting their bodies from the pitch angle used during the strike to a more neutral horizontal position. This repositioning takes time and may represent a trade-off between achieving dual jaw contact, which may be important for feeding performance, and the time and energy expended on repositioning between bites. Body orientation of a prey fish relative to its predator is also known to impact the escape probability of the prey (Kimura & Kawabata, 2018), suggesting that the relationship between body pitch and jaw contact may have implications for antipredator behaviors and escape probabilities among fishes feeding on attached prey. It is also possible that shallower body angles permit *N. nematopus* to expend less energy during grazing by presenting a smaller profile compared to grazing with a steeper body angle, which may be important in environments with high water flow. Previous work characterizes *N. nematopus* as having a postcranial morphological profile adapted for lotic, as opposed to lentic, habitats, providing support for this hypothesis (Říčan et al., 2016).

We also note that *N. nematopus* occupies an extreme position along the primary axis of variation for the combined 9-variable dataset and is set apart by the first and second principal components of kinematic and behavioral variation, respectively. *Neetroplus nematopus* is further distinguished from the other six species by its lateral body pitch angle, which is the lowest and least variable in the dataset, its rate of dual-jaw contact, which is the highest in the dataset, and its unique ability to maintain dual-jaw contact at relatively shallow body angles. This may reflect greater specialization for benthic-feeding in this species compared to *H. multispinosa* or *A. nigrofasciata*, and may confer performance advantages compared to other less specialized benthic-feeding species. Further work is needed to test this hypothesis.

## ACKNOWLEDGEMENTS

We thank Kirk Winemiller for his insights regarding the diet of *Kronoheros umbriferus,* Michalis Mihalitsis, who helped develop the experimental food recipe, and Jasen Liu, who provided valuable feedback on the manuscript. Finally, we are grateful for assistance with film collection, frame extraction, and landmark placement from Chloe Kaelin, Lauren Friker, Joey Kennedy-Van Gilst, Ivan Ross, Claire Low, and Ashley Zapanta.

## AUTHOR CONTRIBUTIONS

PCW and KTR conceptualized the project. KTR procured specimens, collected primary data, and analyzed data. PCW and KTR interpreted data. KTR prepared and wrote the manuscript. PCW and KTR edited the manuscript. PCW provided funding for project materials.

## DATA & RESOURCE AVAILABILITY

All data files used in this study are available from the first author. We used the following software: RStudio (source: https://posit.co/download/rstudio-desktop/), ImageJ (source: https://imagej.net/ij/download.html), tpsDig (source: https://www.sbmorphometrics.org/soft-dataacq.html).

